# mRNA-LNP Vaccines Encoding for Dengue Optimized prM/ENV Proteins Induce Protective Immunity without ADE

**DOI:** 10.1101/2025.07.21.665855

**Authors:** Enzo LaMontia-Hankin, Clayton J Wollner, E Taylor Stone, Angélica Peña Rosado, Jill Pflugheber, Ashley Zalla, Sebastian Nieto, Amelia K Pinto, James D Brien, Justin M Richner

**Affiliations:** Department of Microbiology and Immunology, University of Illinois College of Medicine, Chicago, Illinois, USA; Department of Microbiology, Immunology & Molecular Genetics, University of Kentucky, Lexington, Kentucky, USA

## Abstract

Dengue virus (DENV) is the most common mosquito-borne virus in the world, causing nearly 400 million infections annually, with the number of cases predict to increase over time. Despite this prevalence there are no widely approved vaccine for DENV naïve individuals or therapeutics. DENV consists of four distinct serotypes, DENV 1-4, that share 60-70% homology. Development of a safe and efficacious pan-Dengue vaccine has been complicated by the potential of antibody-dependent enhancement, in which cross-reactive non-neutralizing antibodies can enhance infection and disease. We previously demonstrated that a mRNA-LNP encoding for DENV-1 prM and Envelope proteins generates homotypic neutralizing immune responses and protects against a lethal challenge. The DENV-1 vaccine avoided ADE by eliminating the fusion loop (FL), one of the dominant ADE epitopes. Using a similar design methodology, here we present mRNA-LNP prM-E vaccines for DENV 2, 3, and 4. These vaccines maintained the ΔFL modifications, but encode a chimeric E protein to improve VLP expression and stability. All three vaccines elicited neutralizing titers of serotype specific antibodies mice and protected against a lethal homotypic DENV challenge. Mutation of the FL lowered ADE in vaccinated mouse sera. We formulated monotypic vaccines into a tetravalent vaccine and evaluated for immunogenicity and protection in mice. The tetravalent vaccine elicited neutralizing humoral responses against all DENV serotypes and protected against lethal challenges. Cumulatively, our monovalent DENV mRNA-LNP vaccines against all DENV serotypes generate protective immunity and lower the potential for ADE, leading to the development of a first-generation ADE-altered tetravalent DENV vaccine.

## INTRODUCTION

Dengue virus (DENV) is the most common vector-borne virus to infect humans, infecting an estimated 100-400 million people annually.^1^ DENV is a member of the family *Flaviviridae*, where it is most closely related to Zika virus (ZIKV) and more distantly related to West Nile virus (WNV), Yellow Fever virus (YFV), and Japanese Encephalitis virus (JEV).^6^ DENV consists of 4 distinct serotypes, DENV 1, 2, 3, and 4, which can cause similar disease outcomes.^7^ While these serotypes share 60-70% sequence homology, they are mostly antigenically unique, with some cross-reactive epitopes.^1^ DENV’s ∼11kb positive sense ssRNA genome encodes a single polypeptide that contains three structural proteins and seven nonstructural proteins.^1–3^ The structural proteins premembrane (prM) and envelope (E) comprise the virion and are critical to the antigenic landscape, while the capsid (C) binds and stabilizes the viral genome.^4^ The primary DENV vector are *Aedes* genus mosquitos, predominately *Aedes aegypti*, and to a lesser degree *Aedes albopictus*.^1^ Approximately 3.9 billion people throughout Central/South America, Asia, and various Pacific islands currently live in the range of DENV’s vectors, posing an enormous public health threat.^1,5^ DENV is a member of the family *Flaviviridae*, where it is most closely related to Zika virus (ZIKV) and more distantly related to West Nile virus (WNV), Yellow Fever virus (YFV), and Japanese Encephalitis virus (JEV).^6^ DENV consists of 4 distinct serotypes, DENV 1, 2, 3, and 4, which can cause similar disease outcomes.^7^ While these serotypes share 60-70% sequence homology, they are mostly antigenically unique, with some cross-reactive epitopes.^1^

The majority of DENV infections are asymptomatic, but the most common symptomatic disease outcome is Dengue Fever (DF).^8^ DF consists of severe widespread pain as well as high fevers (>103°F/40°C), rashes, and nausea.^1,2^ DF symptoms typically resolve 1-2 weeks after onset. In rare cases, DF can progress into more severe Dengue Hemorrhagic Fever (DHF) and Dengue Shock Syndrome (DSS)^2^, with the additional symptoms of vascular leakage, resulting in bleeding in mucosal membranes, urine, and stool.^1,8,9^ Without medical intervention, untreated DHF/DSS can be fatal via hypovolemic shock and organ failure. Currently there are no approved therapeutics and no widely approved vaccines for DENV, with the current standard of care of DHF/DSS patients to be hospitalization with IV fluids and pain management.^2,8^

Typically, primary infection of any DENV serotype results in long term protection against the infecting serotype, but not the other three serotypes.^10–12^ Despite this long-lived protection, neutralizing antibodies comprise a minority of the total polyclonal response. The majority of the antibodies in the polyclonal response target cross-reactive and poorly/non-neutralizing epitopes on immature prM or the fusion loop (FL) within domain II of the E protein.^11,12^ These antibodies can drive antibody dependent enhancement (ADE) in heterotypic secondary infections, in which case the infecting virus is bound by non-neutralizing antibodies from the primary infection, leading to FcγR-mediated phagocytosis. Once the opsonized-DENV enters the endosome of the FcγR+ cell, DENV E proteins undergo conformational changes that reduce efficacy of bound antibodies and induce viral fusion in the acidified endosome, subsequently infecting the cell.^13–16^ Because ADE is significantly associated with more severe DENV disease outcomes, it is critical to evaluate any DENV vaccine formulations’ capacity to generate enhancing antibodies.^14^

Because of the potential of ADE, the antibody epitopes presented by within a DENV vaccine must be selected carefully. Antibodies specific to tertiary and quaternary epitopes, which are only found on intact viral particles, are necessary for neutralizing antibody responses.^12,17^ To prevent ADE, a DENV vaccine must elicit a balanced, neutralizing response to all serotypes targeting neutralizing epitopes and not ADE-enhancing epitopes. The first widely approved DENV vaccine, Dengvaxia, was a tetravalent live attenuated vaccine (LAV) based off the YFV-17D vaccine.^18^ While Dengvaxia was protective in seropositive individuals, it failed to drive a long lasting and protective immune response to all DENV serotypes in DENV-naïve individuals.^19,20^ Because of this, seronegative children who received Dengvaxia were at an increased risk of severe disease or death during a primary infection due to ADE.^18,21^ Because of this increased risk of severe disease, all future DENV vaccine candidates must be designed to mitigate ADE, while maintaining relevant neutralizing epitopes.

The mRNA vaccine platform has proven to be safe and efficacious with billions of doses delivered worldwide during the COVID-19 pandemic. For flaviviruses, the RNA platform represents a major advantage over the leading LAV platforms. All leading DENV LAV’s include an unmodified FL to support viral replication. With the mRNA-LNP platform we can engineer mutations into the coding RNA at the FL in order to direct the host immune response away from ADE-driving epitopes. Unlike other viruses, such as SARS-CoV-2 and the spike protein, where expression of a single structural protein is adequate to elicit a neutralizing humoral response, recombinant DENV E (rE) likely lacks the most important neutralizing epitopes. rE subunit vaccines drive inadequate immunogenicity in preclinical settings likely due to structural issues and a lack of quaternary epitopes.^22–24^ Upon translation of the prM/ENV polyprotein from ribosomes in the host ER, host proteases cleave the prM-E and these viral proteins are folded and anchored in the ER membrane. Appropriate folding and trafficking of the viral proteins is aided by a signal peptide at the N-terminus of the polyprotein. Expression of prM and E results in the de novo formation of a non-replicative viral like particle (VLP) with a T=1 symmetry.^25^ VLPs preserve the majority of highly neutralizing quaternary epitopes, which direct the host immune response towards neutralizing B and T cell epitopes.^25–27^

We first developed a novel mRNA-LNP DENV 1 vaccine that achieves high levels of *in vitro* VLP expression. *In vivo*, the vaccine elicited high levels of protective and non-cross-reactive antibodies in an immunocompetent C57BL/6 mouse model, and was 100% protective in immunocompromised AG129 mice.^28^ Here, we have developed three novel monovalent prM-E mRNA-LNP vaccines for DENV 2, 3, and 4 based on our original DENV 1 vaccine. Each monovalent vaccine encodes prM and modified E of a given serotype. These monovalent vaccines showed strong neutralizing antibody responses in WT C57/BL6 mice, while eliminating or reducing ADE. All monovalent vaccines protected mice from a lethal homotypic challenge. We then combined the DENV 1, 2, 3, and 4 monovalent mRNA-LNPs into a tetravalent vaccine. Tetravalent vaccines drove a mixed response against the four DENV serotypes with protective immunity against DENV 3 and DENV 4. These data indicate a significant step towards an effective and safe tetravalent DENV vaccine.

## RESULTS

### Design of DENV 1, 2, 3, and 4 prM/E constructs and *in vitro* expression

Similar to our previously published DENV 1 approach, we designed nucleotide sequences for dengue serotypes 2, 3, and 4 containing prM and E chimeric proteins from DENV 2 (strain Thailand 16684/84), DENV 3 (strain Sri Lanka D3/H/IMISSA/SRI/2000/1269), and DENV 4 (strain Dominica/ 814669/1981).^28^ Similar to previous observations, we found that robust expression and VLP secretion of the prM/ENV antigens of DENV2, 3, and 4 required optimization of the stem/transmembrane domain of ENV.^25,29,30^ The native stem and transmembrane domains of each E protein was replaced with the sequence from JEV.^31^ Lastly, variants of the DENV 2, 3, and 4 sequences were modified to include 3 mutations (G106R, L107D, and F108A) in the fusion loop to ablate the epitope, which has been shown to reduce the generation of poorly neutralizing but cross-reactive antibodies associated with ADE.^28,30^ Directly upstream of the prM and E sequence, we included the sequence for the tissue plasminogen activator (tPa) signal peptide which we have shown previously to increase VLP secretion with our DENV 1 vaccine.^28^ All sequences contain a 5’ UTR and 3’ UTRs that have been used in various mRNA vaccines.^26,28,32–34^

All nucleotide sequences were *in vitro* transcribed via a T7 RNA polymerase promoter site directly upstream from the 5’ UTR. To camouflage our vaccine RNA from host innate RNA sensors, we include the modified ribonucleotide N1M-Ψ Uridine in lieu of standard Uridine.^35–37^ *In vitro* transcription produces double stranded RNA artifacts that can activate intracellular innate immune sensors that are known to reduce translation of exogenous mRNA. Double-stranded RNA contamination was removed from IVT-synthesized RNA thought a cellulose purification step.^38^ Finally, a 5’ cap-1 structure and 3’ poly(A) tails were added enzymatically to yield RNA that mimics host mRNA (Fig 1A). These mRNAs were encapsulated into lipid nanoparticles using either Genvoy (DENV 2) or SM-102 (DENV 3/4) ionizable lipids. To confirm VLP expression *in vitro*, HEK293T cells were transfected with vaccine RNA and supernatants were harvested 48 hours post transfection. Purified VLP’s were then imaged with a transmission electron microscope, and particles of approximately 30nm were detected with all constructs (Fig. 1B). While DENV virions are ∼50nm, it has been reported similar antigenicity between DENV VLP’s and virions.^25^ Thus, deletion of the fusion loop did not inhibit VLP formation and secretion, similar to our previous results with DENV 1 ΔFL prM/E mRNA vaccines. To evaluate the *in vivo* humoral response, 20µg of the mRNA-LNP vaccines were evaluated identically as described above (Fig 1C). Mice received the boost dose 28 days post prime, and animals were sacrificed 28 days post boost.

**Figure 1:**
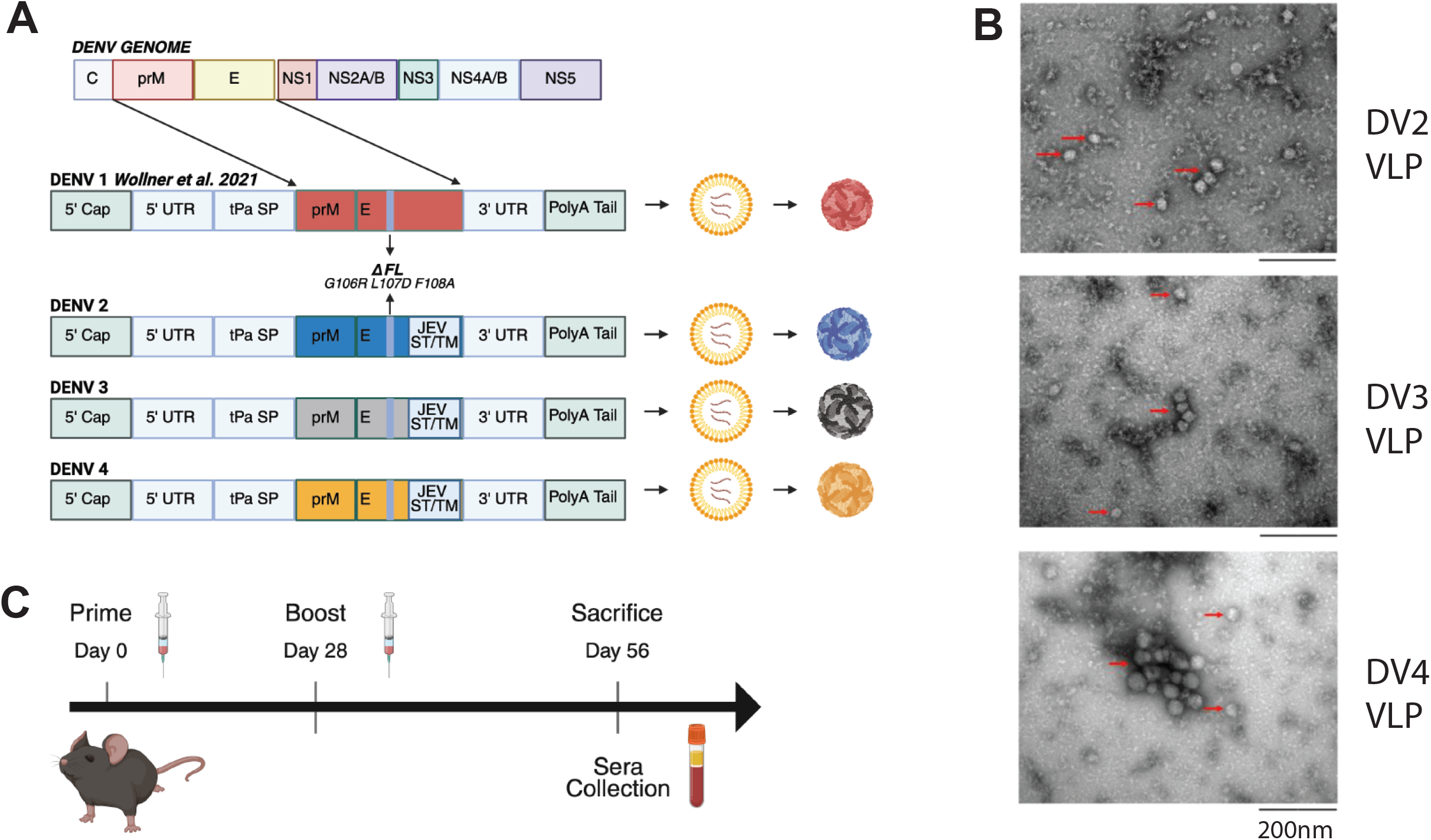
DENV prM/E vaccine design, viral protein expression, and *in vivo* vaccination scheme. (A) DENV genome and schematic showing previously published and current vaccine design. Four separate mRNA’s were engineered to encode prM and E proteins from a single DENV serotype. (B) TEM images of extracellular VLP’s. HEK 293T cells were transfected with each vaccine mRNA and supernatant was harvested. Dialyzed supernatants were imaged via TEM. (C) Vaccine schedule used for all *in vivo* analysis. C57BL/6 were vaccinated on a two-dose prime-boost schedule intramuscularly.

### DENV 2 prM/E Vaccine adaptive immune response

C57BL/6 mice were vaccinated as described above. We included a group of mice that received a mRNA-LNP encoding GFP as a negative control and a positive control cohort of mice that were infected with 1E5 focus forming units (FFU’s) of DENV 2 (strain New Guinea C, NGC) to compare the humoral response of the vaccine to that of a DENV 2 infection in an immunocompetent mouse. Total DENV specific IgG from heat inactivated vaccinated mouse sera was quantified via endpoint dilution ELISA against purified DENV 2. All ΔFL vaccinated mice and four out of five WT vaccinated mice seroconverted and had significantly higher DENV 2 specific IgG levels than GFP vaccinated mice (Fig. 2A). DENV 2 WT vaccinated animals had an average endpoint dilution of ∼1/33,000 and DENV 2 ΔFL vaccinated animals had an average endpoint dilution of ∼1/55,000 compared to the GFP control of ∼1/3,400 (*P* values = .0065 and .0013 respectively). DENV specific IgG titers from both vaccine variants were not significantly different compared to DENV 2 infected animals (*P* values = .5887 and .9680 respectively). These data indicate the mRNA vaccines elicit a humoral response of similar magnitude to a natural infection.

**Figure 2:**
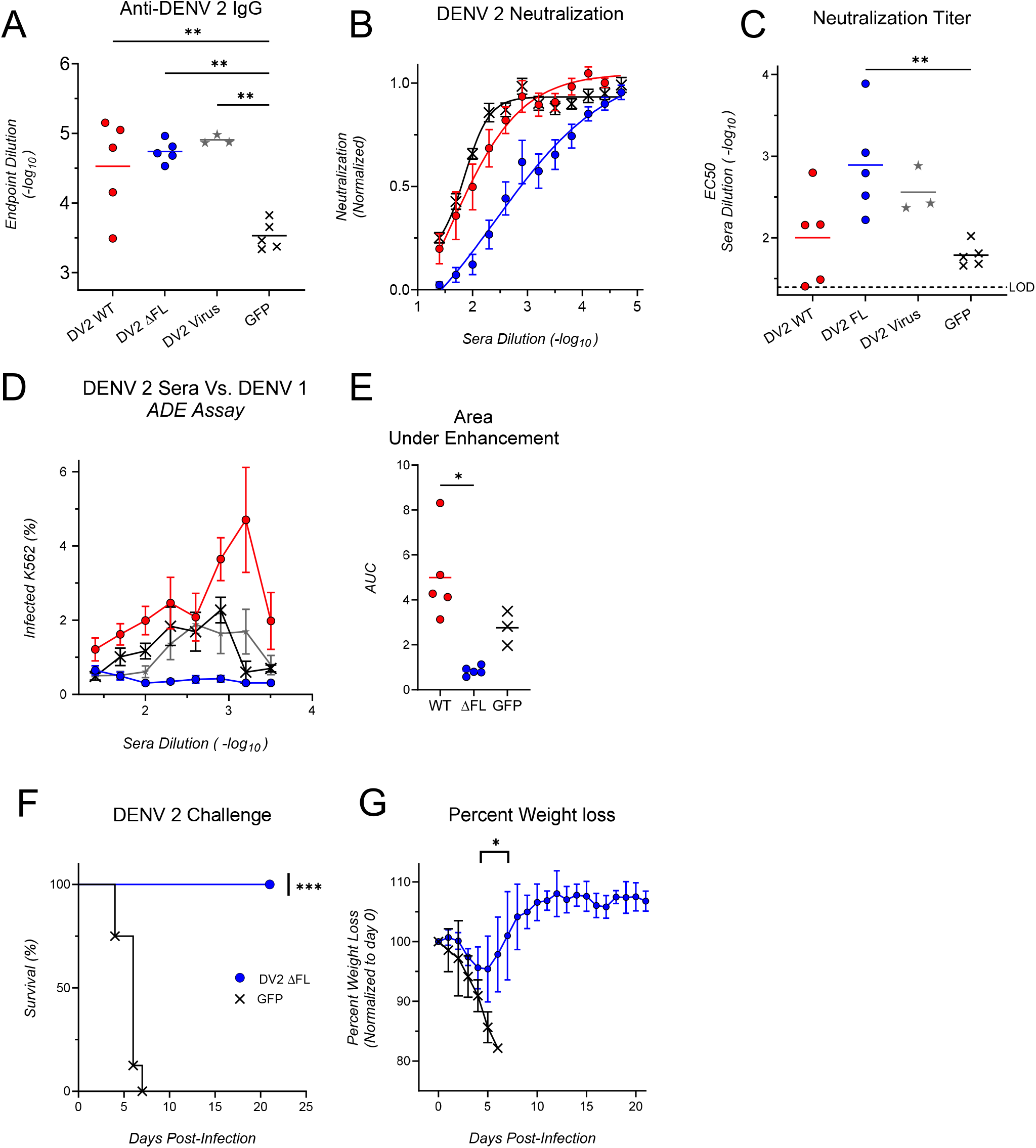
DENV 2 prM/E ΔFL vaccine induces neutralizing, non-enhancing antibody responses. (A-F) C57BL/6 mice were vaccinated with the WT or ΔFL DENV 2 vaccine or a GFP control (N=5/group). Another group of mice (N=3) were infected with 1E5 FFU’s of DENV 2 NGC. (A) Serial dilutions of sera was analyzed for binding of purified DV2 via ELISA assay. The endpoint dilution was determined for each animal. (B) Focus reduction neutralization tests (FRNT) were conducted with serial dilutions of sera against DENV 2. Each point represents the mean of the experimental group +/- the standard error of the mean (SEM) (C) EC50’s from neutralization curves of individual vaccinated mice are reported. (D) Serial dilutions of vaccinated animal sera were analyzed for enhancement of heterologous DENV 1 16007. (E) The area under the enhancement curves for each individual mouse is also reported. F-G) IFNAR^-/-^ were vaccinated with 10µg of the ΔFL DENV 2 vaccine or a GFP encoding mRNA-LNP. Animals were challenged with1x10^4 FFU of DENV 2. Survival curve (F) and weight loss (G) was reported. *, P < 0.05; **, P < 0.01; ***, P < 0.001.

The neutralizing antibody titer was determined by Focus Reduction Neutralization Test (FRNT) against DENV 2 NGC (Fig 2B). The serum concentration at which 50% viral neutralization occurs was reported as EC50 for individual mice (Fig 2C). Despite significant DENV 2 specific IgG the WT vaccine failed to drive a significant neutralizing response, with a mean EC50 of ∼1/100 and a maximum EC50 of ∼1/626. This, compared to the mean GFP EC50 of ∼1/60 was insignificant (*P* value = .82). However, the ΔFL vaccine had a mean EC50 of ∼1/780 with a maximum EC50 of ∼1/7,700 compared to GFP vaccinated control (*P* value = .0068). Neutralization titers were equivalent between the ΔFL vaccinated mice and the naturally infected mice, again indicating a humoral response of similar magnitude.

In a natural DENV infection, cross-reactive heterotypic antibodies comprise the majority of the polyclonal response, which can cause ADE.^12^ We characterized the ADE capacity of both variants of our DENV 2 vaccines. We quantified ADE by performing serial dilutions of our DENV 2 vaccine and opsonizing those dilutions with a constant MOI of DENV-1 virus. While sera is collected 28 days post boost, which is not enough time for neutralizing antibodies to wain for ADE to occur, we do serial dilutions of the sera to simulate waning humoral immunity. We then infected FcγR+ K562 cells with the sera opsonized DENV-1. The frequency of infected cells was determined via flow cytometry using intracellular staining with 4G2, a pan-DENV monoclonal antibody specific to the FL. Sera from WT DENV 2 vaccinated mice elicited a ∼1.7 fold higher infection rate compared to virus only controls, indicating a modest enhancement. Peak infection with the ΔFL DENV 2 vaccine was ∼4.2 fold lower than virus only infection, indicating a lack of enhancement (Fig. 2D). ADE of the individual mice are plotted as area under the curve. The sera from WT vaccinated caused significantly more ADE compared to sera from the ΔFL vaccinated mice (Fig 2E, p-value = 0.002)

To understand the protective capacity of the DENV 2 vaccines, a separate study was conducted with susceptible mice which lack the which lack the type I IFN receptor (IFNAR^-/-^). In the same vaccine administration schedule, IFNAR^-/-^ mice were given 10 µg’s of the DENV2 ΔFL or the mock GFP vaccine. Vaccinated IFNAR^-/-^ mice were challenged with 1x10^4^ FFU of DENV 2 strain D2S20. All DENV 2 vaccinated mice survived the challenge, while all GFP vaccinated animals died by day 7 (*P* value = .0003) (Fig. 2F). Minor weight loss was observed with DENV-2 vaccinated animals, with a ∼5% mean weight loss occurring at day 5 (Fig 2G). By day 7, the mean weights returned to day 0 levels, and by day 9 all animals had fully returned to day 0 weight. These data demonstrate that the DV2 prM/ENV mRNA-LNP vaccines elicit protective immune responses.

### DENV 3 prM/E Vaccine adaptive immune response

*In vivo* immunogenicity assays of the DENV 3 mRNA-LNP vaccine were conducted as previously described (Fig 1C). Both WT and ΔFL variants of the DENV 3 vaccine were used at 20µg doses in a prime-boost schedule. While infection with DENV 2 NGC elicits a humoral response, similar infection with DENV 3 H87 failed to drive a detectable humoral response in WT mice, so no viral infection comparator control was included. Animals were sacrificed 56 days post prime, and sera was collected for humoral analysis. To quantify the DENV 3-specific IgG response, heat inactivated sera from vaccinated animals was quantified via endpoint dilution ELISA coated with purified DENV 3. All vaccinated animals seroconverted and had statistically significant DENV 3 specific IgG’s compared to a naïve control. The mean endpoint dilution of both WT and ΔFL variants were 1/213,484 and 1/205,966 respectively which was significantly elevated compared to the endpoint of naïve control sera at 1/1,268 (*P* values = <0.0001) (Fig. 3A). To quantify the neutralizing antibody response, heat-inactivated sera was analyzed via homotypic FRNT. Compared to the naïve cohort (EC50= 1/26), both WT and ΔFL cohorts had significantly elevated homotypic neutralizing antibody titers, with the average EC50 of ∼1/337 for the WT cohort and ∼1/269 for the ΔFL cohort (*P* value = <.0001 and .0002 respectively) (Fig. 3B/C).

**Figure 3:**
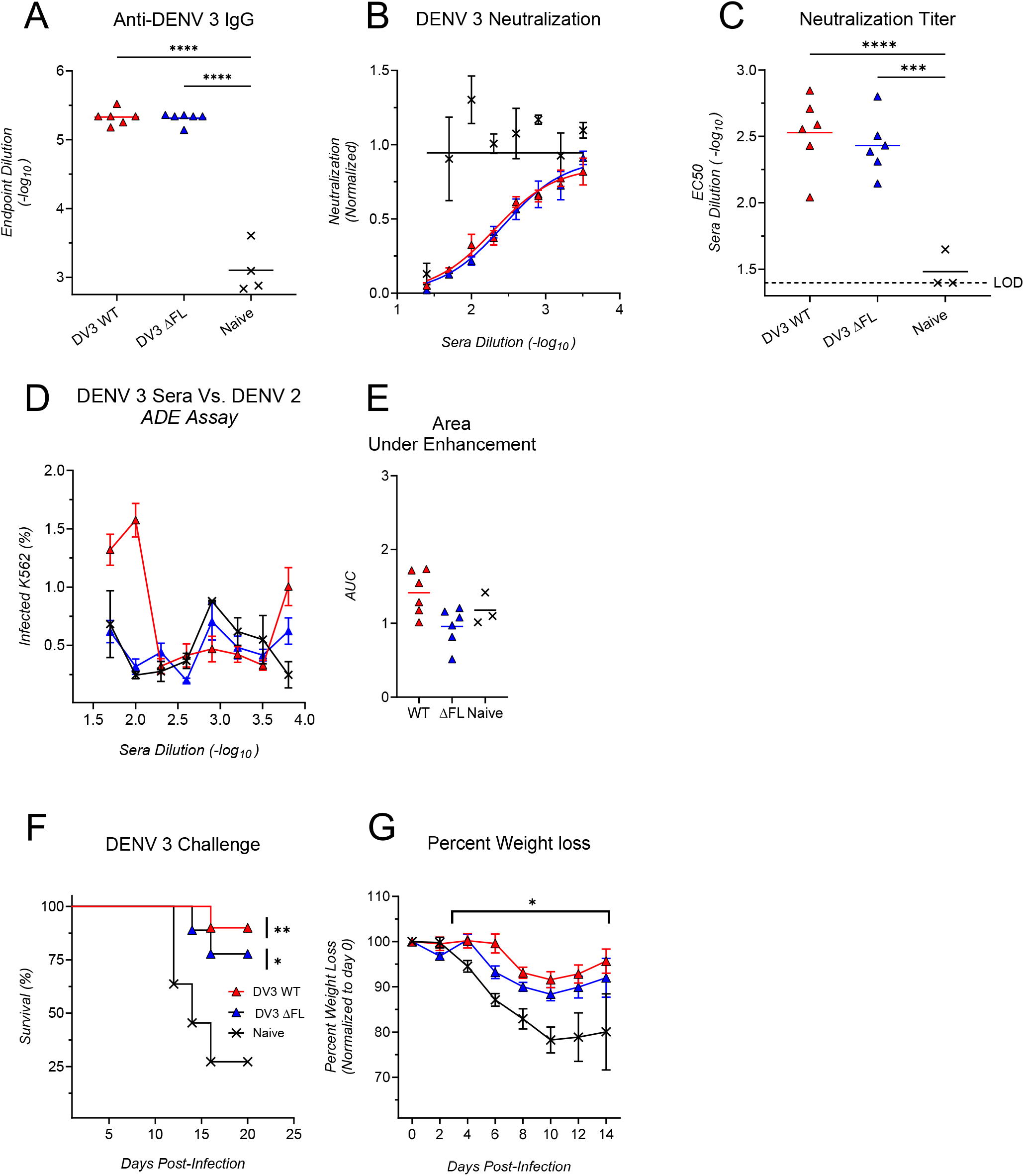
DENV 3 prM/E vaccines induces neutralizing, non-enhancing antibody responses. Immunocompetent C56BL/6 mice (N=6) were vaccinated prime-boost with either 20µg of the WT or ΔFL DENV 3 vaccine. (A) Serial dilutions of heat inactivated vaccinated mouse sera was applied to an ELISA plate coated with purified DENV 3 to quantify total antiviral IgG. The endpoint dilution of each mouse is plotted. (B) Focus reduction neutralization tests (FRNT) were conducted with serial dilutions of vaccinated heat inactivated animal sera against DENV 3 H87. (C) EC50’s from neutralization curves of individual vaccinated mice is reported. (D) Serial dilutions of vaccinated animal sera were analyzed for heterotypic enhancing antibodies with DENV 2. (E) The area under the enhancement curves for each individual mouse is plotted. (F-G) IFNAR1^-/-^ mice (N=9-11) were vaccinated with 10µg of either the WT or ΔFL DENV 3 vaccine, or a PBS control. Animals were challenged with 1x10^7 FFU of DENV 3. Survival curve (F) and weight loss (G) was reported. *, P < 0.05; **, P < 0.01; ***, P < 0.001; ****, P < 0.0001.

DENV 3 vaccinated sera was also analyzed via ADE assays against heterotypic DENV 2 (Fig 3D/E). Sera from WT vaccinated animals showed a modestly enhanced peak infection of (∼1.6% compared to the ∼1.45% DENV-2 only infection) while the ΔFL vaccinated sera had a peak infection of .7%. These data indicate that antibodies from the DENV 3 ΔFL vaccine do not drive heterotypic enhancement with DENV 2.

To investigate the protective capacity of the DENV-3 vaccines, IFNAR^-/-^ animals were vaccinated in a prime-boost manner with 10µg of the WT or ΔFL variants, or mock vaccinated controls. Mice were challenged with 1x10^7^ FFU DENV 3 C0360/94. Both vaccine variants were significantly protective from a lethal DENV 3 challenge compared to a mock vaccinated control. While both variants were not 100% protective, the WT vaccine yielded a 90% survival rate (*P* value = .0029) and the ΔFL vaccine yielding a 77% survival rate (*P* value = .0204) (Fig. 3F). The vaccinated animals also had significantly lower weight loss compared to mock vaccinated animals (Fig 3G). Both DENV 3 vaccines elicit neutralizing, serotype specific, and protective humoral immunity.

### DENV 4 prM/E Vaccine Adaptive immune response

In vivo evaluation of the DENV-4 vaccines followed an identical vaccination scheme used with the DENV 2 and DENV 3 vaccines (Fig 1A). C57BL/6 mice were vaccinated with 20µg of the WT or ΔFL variant of the mRNA-LNP vaccine in a prime-boost manner. Similar to DENV 3 H87, DENV 4 UIS 497 did not elicit a humoral response in immunocompetent mice and thus was omitted from this analysis. 56 days post prime animals were sacrificed, and sera was collected for analysis. To quantify the total DENV 4 specific IgG response to vaccination, heat inactivated sera was analyzed via endpoint dilution ELISA against purified DENV 4. All ΔFL DENV 4 vaccinated animals and 3/5 WT DENV4 vaccinated animals seroconverted (Fig. 4A). Mean endpoint dilutions for the WT and ΔFL variants were significant compared to naïve control at ∼1/47,785 and ∼1/101,728 respectively (Fig 4B, *P* value = .0042 and 0005 respectively).

**Figure 4:**
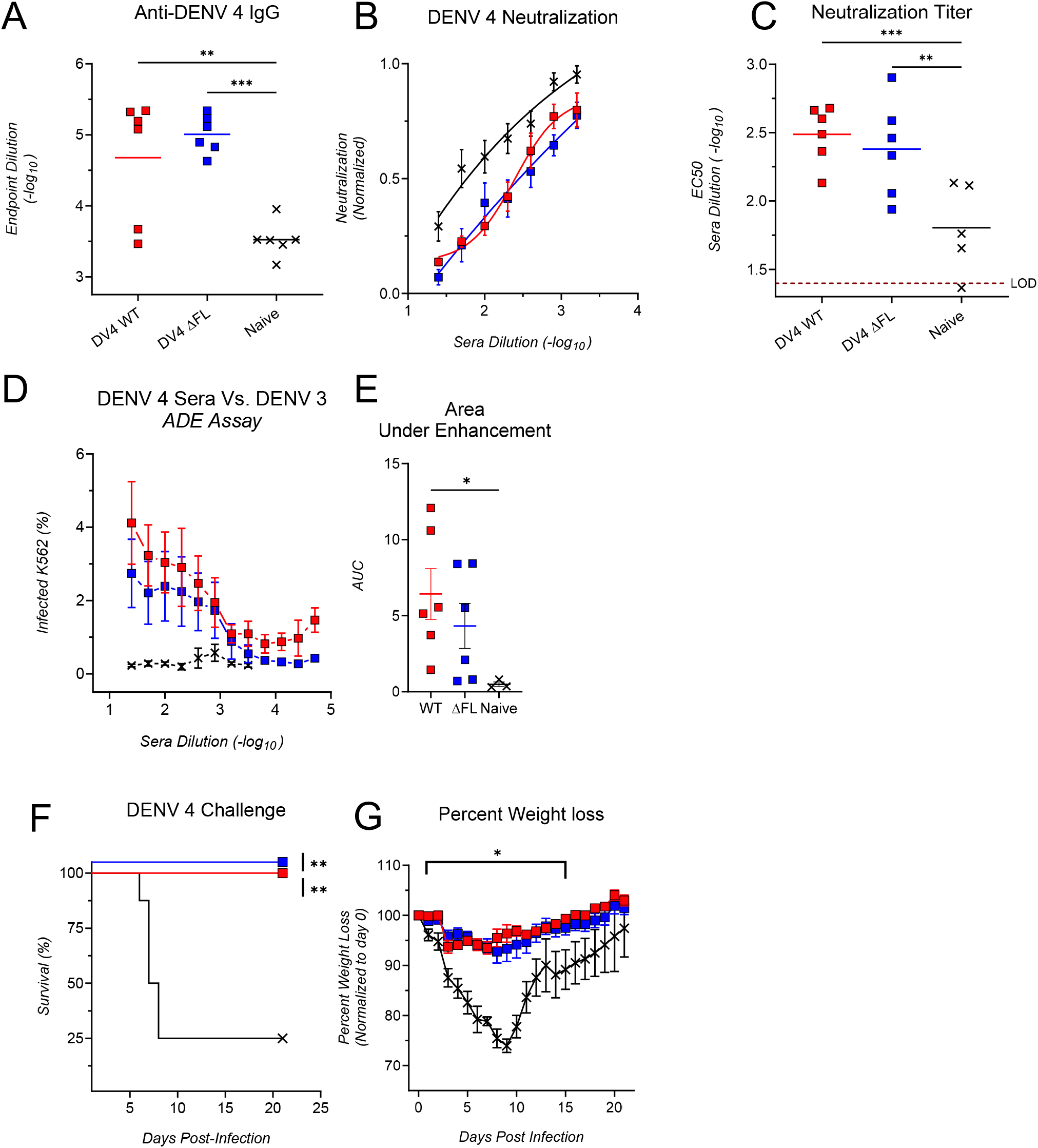
DENV 4 prM/E vaccines induces neutralizing, non-enhancing antibody responses. C57BL/6 mice (N=6/group) were vaccinated with either 20µg of the WT or ΔFL DENV 4 vaccine. (A) Serial dilutions were analyzed for total anti-DENV 4 IgG via ELISA assay and endpoint dilution of individual mice determined. (B) FRNT were conducted with serial dilutions of vaccinated sera against DENV 4. (C) EC50’s from neutralization curves of individual vaccinated mice is reported. (D) Sera were analyzed for heterotypic enhancing antibodies with DENV 3. (E) Area under the curve of individual mice are reported. (F-G) IFNAR1^-/-^ mice (N=8-10/group) were vaccinated with 10µg of either the WT or ΔFL DENV 4 vaccine, or a PBS control. Animals were challenged with 5x10^6 FFU of DENV 4. Survival curve (F) and weight loss (G) was reported. *, P < 0.05; **, P < 0.01; ***, P < 0.001.

Sera neutralization was also analyzed via FRNT with a homotypic DENV-4 strain. Despite the ΔFL vaccine yielding ∼2X higher DENV specific IgG, the ΔFL had a slightly lower average EC50 compared to the WT variant (Fig. 4B/C). The WT variant yielded a mean EC50 of ∼1/307, while the ΔFL variant had a mean of ∼1/240. Despite the marginally lower mean, the peak EC50 for the ΔFL variant was ∼1/780, while the peak EC50 for the WT variant was ∼1/477. The EC50 for both variants’ EC50’s was significantly elevated compared to naïve control, indicating both vaccine variants induced neutralizing humoral responses (*P* value = .001 and .006 respectively).

Antibody cross-enhancement was evaluated via ADE assay, using DENV 3 virus. Both WT and ΔFL showed increased enhancement compared to naïve sera. Notably, the WT vaccine elicited higher, but statistically insignificant, mean enhancement compared to the ΔFL variant, indicating the ΔFL mutations had a modest effect at reducing cross reactive antibodies (Fig 4D). When looking at the area under the curve of individual mice (Fig 4E), both vaccine groups had considerable heterogeneity in their cross-reactive enhancement response.

To investigate the protective capacity of the DENV 4 vaccines, IFNAR^-/-^ animals were vaccinated in a prime-boost manner with 10µg of the WT or ΔFL variants, or mock vaccinated controls. Mice were challenged with 5x10^6^ FFU DENV 4 TVP376. Both vaccine variants completely protected against a DENV-4 challenge. Mock vaccinated control mice had 75% lethality, P value < 0.001 (Fig. 4F). The vaccinated animals also had significantly lower weight loss compared to mock vaccinated animals (Fig 4G). Both DENV-3 vaccines elicit neutralizing, serotype specific, and protective humoral immunity.

### Tetravalent Vaccination Adaptive Immune Response

As we have developed effective monovalent mRNA-LNP vaccines for all DENV serotypes, we now sought to characterize the vaccines in a tetravalent delivery setting. To characterize the protective capacity of the tetravalent vaccine, we vaccinated 18 IFNAR^-/-^ mice with all four ΔFL vaccine variants with 10µg’s of the DENV 1 vaccine and 20µg’s each of the DENV 2, 3, and 4 vaccines. Animals were vaccinated on an identical prime-boost timeline as previously described (Fig. 5A). Four weeks post boost, animals were bled to characterize pre-challenge humoral responses. The 18 mice were then divided into three groups of 6 and subsequently challenged with lethal doses of either DENV 1, 2, or 4. DENV 3 was omitted from challenge studies due to limitations with DENV 3 lethal challenge stocks.

**Figure 5:**
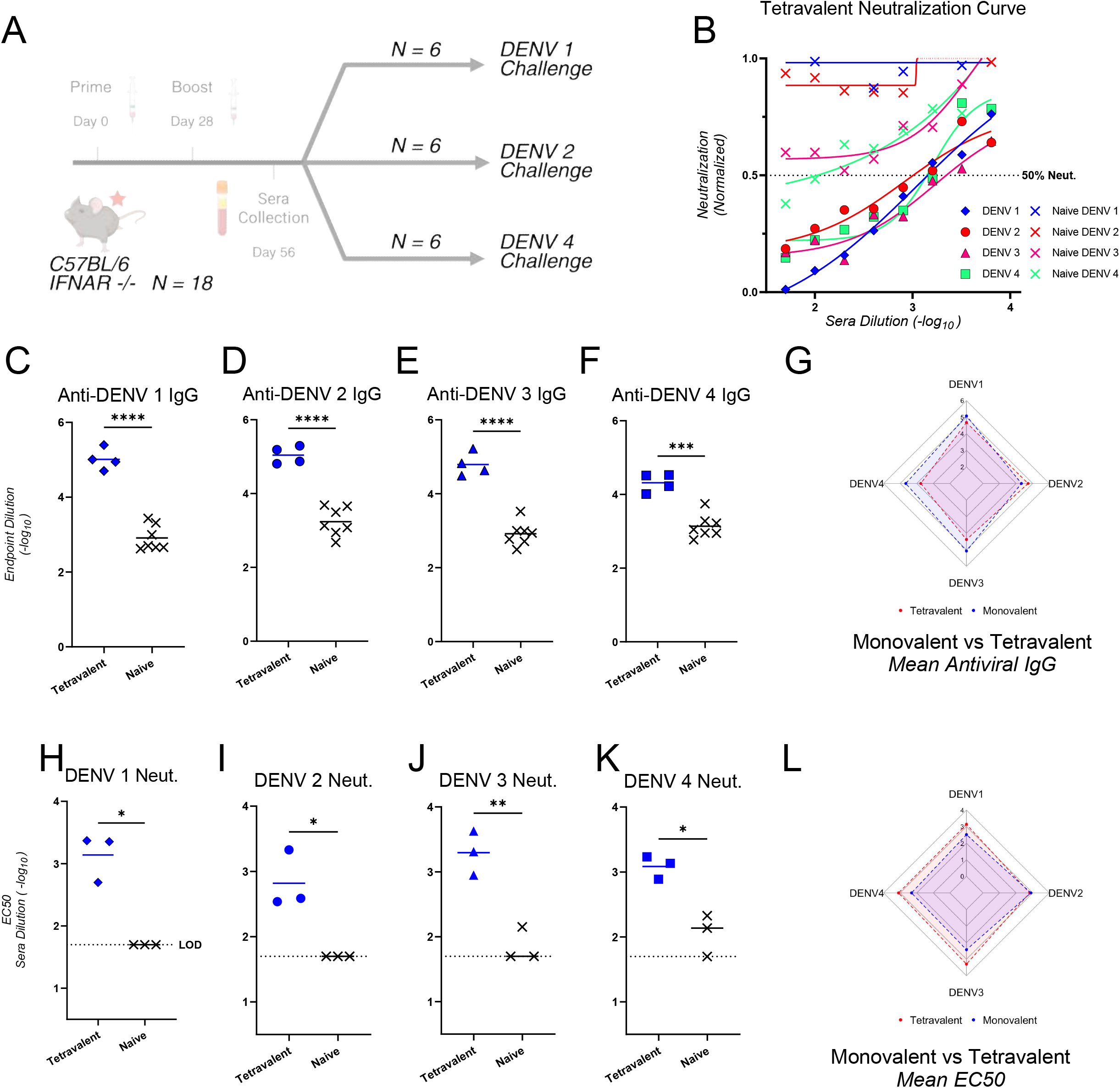
Tetravalent vaccine elicits pan-serotype neutralizing antibody responses. IFNAR-/- mice were injected intramuscularly with ΔFL mRNA-LNP vaccines of each variant in a prime-boost vaccination schedule separated by 4 weeks between the doses. For each dose mice received 10ug DENV1ΔFL and 20ug each of DENV2 ΔFL, DENV3 ΔFL, DENV4 ΔFL mRNA-LNP vaccines. (C-F) Heat inactivated sera was analyzed via ELISA plates coated with a sucrose purified DENV serotype and endpoint dilution is reported. (H-K, B) Sera was serially diluted and analyzed via FRNT with one of four DENV serotypes. EC50’s from neutralization curves are reported. (G, L) Mean EC50’s from experimental groups from monovalent (Fig.’s 2-4) and tetravalent vaccinated mice plotted on a radar graph. *, P < 0.05; **, P < 0.01; ***, P < 0.001; ****, P < 0.0001.

To quantify the anti-DENV IgG, we performed endpoint dilution ELISA’s with heat inactivated pre-challenge sera against all DENV serotypes (Fig. 5C-F). We found significant IgG responses against all DENV serotypes as well as significant neutralizing humoral responses to all DENV serotypes (Fig. 5 H-K). In comparison to the monovalent humoral responses, EC50’s were significantly higher against DENV 1, 3, and 4 in the tetravalent formulation (Fig. 5 L, Table: 1). Despite the increase in neutralizing titer, the type specific IgG was significantly lower against DENV 3 4, with near significance against DENV 1 (P = ∼.08) (Fig. 5 G, Table: 1).

Vaccinated animals were challenged with either 5x10^6^ DENV 1 West Pac, 1x10^4^ FFU’s of DENV 2 D2S20, or 5x10^6^ FFU DENV 4 TVP376. We found the tetravalent vaccine to be 100% protective against lethal DENV 1, 2, and 4 challenges (Fig. 6 A-C). All mock vaccinated animals died, with 100% mortality occurring at day 18 with DENV 1, day 5 with DENV 2, and day 17 with DENV 4 (Fig. 6 A-C). Modest weight loss was observed in the vaccinated, DENV 4 challenged group, with a peak mean weight losses of ∼9% occurring 4 days post infection, with nearly full recovery by day 9 (Fig. 6 F). Weight loss was negligible in vaccinated, DENV 1 and DENV 2 challenged groups (Fig. 6 D, E).

**Figure 6:**
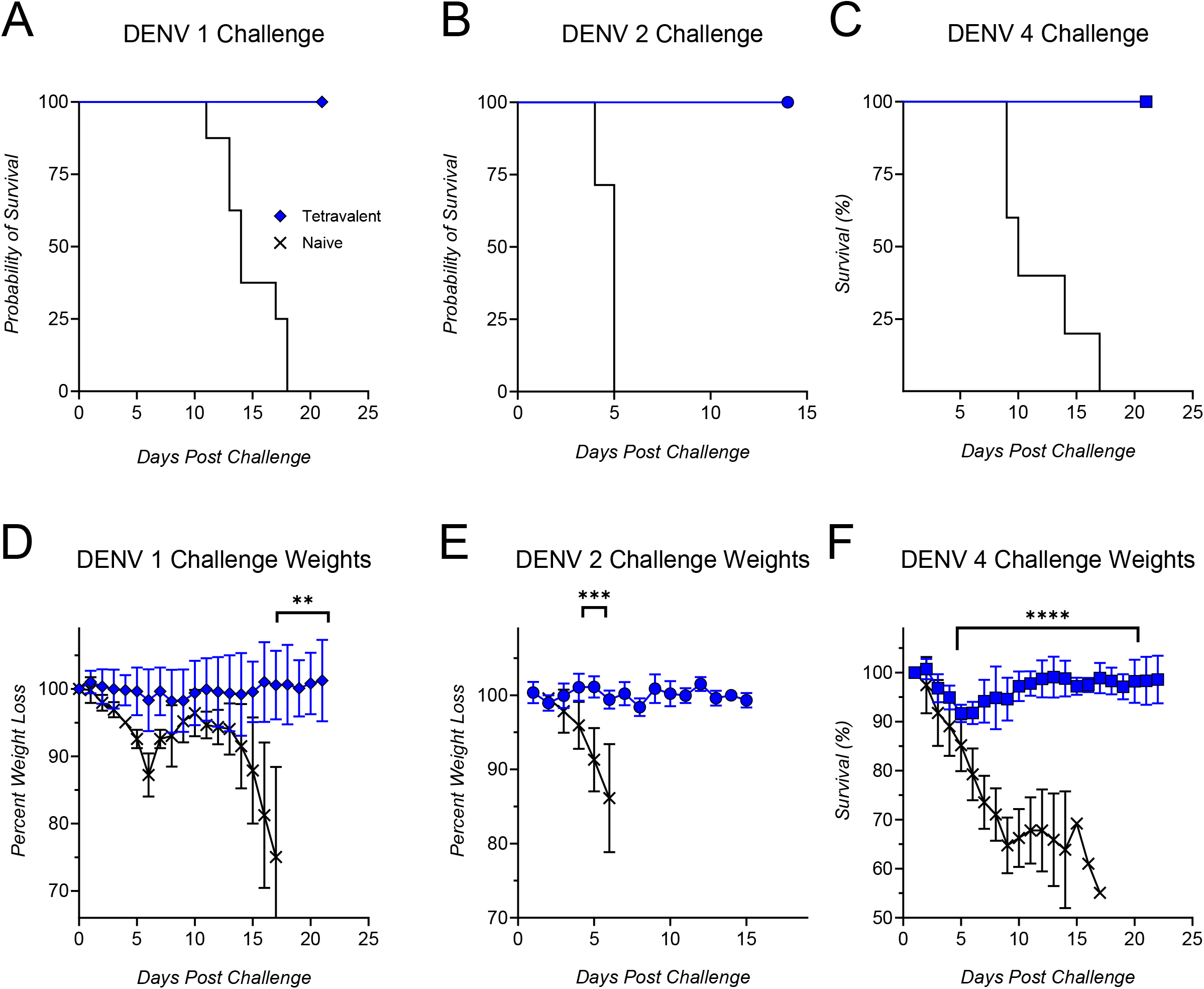
Tetravalent vaccine protects against lethal challenge. (A,D) Six vaccinated mice and 8 naïve IFNAR^-/-^ mice were challenged with 5x10^6 FFU of DENV 1 West Pac. (B,E) Six vaccinated mice and three naïve IFNAR^-/-^ mice were challenged with 1x10^4 FFU of DENV 2 D2S20. (C, F) Six vaccinated mice and five naïve IFNAR^-/-^ were challenged with 5x10^6 FFU DENV 4 TVP376. *, P < 0.05; **, P < 0.01; ***, P < 0.001; ****, P < 0.0001.

## DISCUSSION

While there has been significant innovation and development, there is still no widely approved Dengue vaccine that is safe and broadly efficacious against all four dengue serotypes. DENV vaccine design requires a balance between preserving neutralizing epitopes while minimizing epitopes that drive ADE. The majority of long-lasting humoral immunity is granted by the presence of circulating tertiary and quaternary epitope specific antibodies that are only naturally found on DENV virions.^13,15,39^ This epitope preservation is a major strength of DENV LAV’s, especially considering the generally poor immunogenicity from DENV subunit vaccines.^22–24^ However, because LAV’s are replicative they must also maintain the FL epitope and thus must include epitopes that are known to drive ADE. With DENV, the immunodominant humoral response is weighted heavily towards epitopes on the FL and immature prM.^12,14–16^ If LAV-derived humoral immunity wains to sub-neutralizing levels, vaccine mediated ADE will be possible. With our vaccine design, we are able to preserve the neutralizing epitopes found on LAV’s while also reducing the risk of ADE. Here, along with our previous publication, we have developed nucleotide modified mRNA-LNP vaccine designs for each DENV serotype that drives a protective immune response. In both monovalent and tetravalent settings, our prM-E vaccines drive significant humoral responses to all serotypes, with the ability to protect against lethal challenges.

While the neutralizing humoral response is critical from an efficacy standpoint, it cannot be the only benchmark in which future DENV vaccines are evaluated. The cornerstone of any future DENV vaccine must rest on a deliberate effort to eliminate ADE to make the safest possible vaccine. Emphasizing the safety of any future DENV vaccine is critical in order to regain public confidence given the mistrust generated from Dengvaxia’s initial rollout.^18,21,40^ Crucially, by ablating the FL epitope, we are able to eliminate or reduce ADE *in vitro*. With DENV 2 and DENV 3 vaccinated animals, no ADE was identified with ΔFL variants, and a reduction in ADE was observed with DENV 4. When comparing the AUC’s of these groups, only the WT vs. Naïve areas were significantly different (Fig. 4D). 3/6 ΔFL vaccinated animals had little to no observable ADE, in comparison to 4/6 WT vaccinated animals having a >5% enhancement compared to naïve. While the FL is a significant driver of ADE, other epitopes are known to drive ADE such as immature prM. Elimination of the immature prM epitopes could be achieved by further modification of the coding sequence, including replacement of the native furin cleavage site with a more efficient sequence to drive VLP maturation and hide the immature prM epitope.^41^

Any dengue vaccine must elicit a balanced, neutralizing response against all four serotypes. Notably the monovalent dose required for effective DENV 2, 3, and 4 vaccine-driven immunity was significantly higher than the DV1 vaccine. The prM/ENV Dengue vaccine against DENV 1 yielded robust immunity at low doses of 3μg RNA in a prime/boost immunization strategy in a previous publication from our group.^28^ To achieve similar vaccine-driven immunity, we had to increase the dose to 20μg for DENV 2, 3, and 4 monovalent vaccines. Despite the high dosage, when administered as a tetravalent vaccine, we achieve significant humoral responses against all DENV serotypes. In previous studies, we found that 10µg of the ΔFL DENV 1 vaccine elicited ∼1 log higher neutralizing antibody titers in IFNAR/IFNGR double knock-out mice (AG129) compared to immunocompetent C57BL/6 mice.^28^ Here, we have also found a similar phenotype in our tetravalent vaccine between WT and IFNAR KO C57BL/6 mice, with statistically significant increases in EC50 against DENV 1, 3, and 4, compared to the monovalent setting (Table 1).

**Table 1:** Comparison of total DENV specific IgG and neutralization between monovalent vaccinated and tetravalent vaccinated groups. All mice received the reported vaccine dose, however mouse background differs between monovalent (C57BL/6) and tetravalent (C57BL/6 IFNAR^-/-^).

| DENV-1 ΔFL Vaccine<br>(10μg Dose) | DENV-1 Specific<br>IgG | DENV-1 Neut.<br>(EC50) |
| --- | --- | --- |
| **Monovalent** | 204,000 | 330 |
| **Tetavalent** | 102,000 | 1400 |
| P-Value | NS 0.1692 | * 0.05 |

| DENV-2 ΔFL Vaccine<br>(20μg Dose) | DENV-2 Specific<br>IgG | DENV-2 Neut.<br>(EC50) |
| --- | --- | --- |
| **Monovalent** | 55,000 | 780 |
| **Tetavalent** | 109,000 | 660 |
| P-Value | NS 0.0814 | NS 0.8536 |

| DENV-3 ΔFL Vaccine<br>(20μg Dose) | DENV-3 Specific<br>IgG | DENV-3 Neut.<br>(EC50) |
| --- | --- | --- |
| **Monovalent** | 206,000 | 270 |
| **Tetavalent** | 62,000 | 2000 |
| P-Value | * 0.0421 | * 0.0302 |

| DENV-4 ΔFL Vaccine<br>(20μg Dose) | DENV-4 Specific<br>IgG | DENV-4 Neut.<br>(EC50) |
| --- | --- | --- |
| **Monovalent** | 101,000 | 209 |
| **Tetavalent** | 21,000 | 1300 |
| P-Value | ** 0.0043 | ** 0.0054 |

Interestingly, we observed lower serotype specific IgG titers against DENV 1, 3, and 4 (Table 1), while an increase in the neutralization titers, when comparing tetravalent vaccination in the immunodeficient IFNAR^-/-^ mice compared to the monovalent vaccines in the immunocompetent C57BL/6. These data are surprising given the increased antigenic load and the prevalence for cross-reactive poorly neutralizing antibodies in the tetravalent setting as observed in historical unbalanced immune responses from LAV tetravalent vaccines.^20,42^ With our mRNA-LNP approach, we achieve balanced immunity against all four serotypes counter to previous LAV tetravalent vaccines which drive a dominant response to a single serotype. It is possible that the VLP’s encoded by the mRNA-LNP vaccine are driving a higher quality humoral response by increasing the population of the rare potently neutralizing antibodies that often target tertiary and quaternary epitopes. While our tetravalent vaccine encoded 20µg each of DENV 2, 3, and 4 vaccines, we conducted the monovalent survival studies with 10µg of each vaccine, all of which were significantly protective compared to mock groups. While we were unable to conduct a survival study with tetravalent vaccinated animals against DENV 3, we showed significant DENV 3 specific IgG and neutralization form in these animals, with significantly higher EC50’s compared to monovalent vaccinated animals. We also showed significant protection against a lethal DENV 3 challenge with a 10µg monovalent vaccine, which is half the dose used in the tetravalent formulation.

The “ideal” DENV vaccine strategy has yet to be fully identified by any group, were it so easy. Many studies have shown the need to maintain strongly neutralizing epitopes to drive a durable and protective humoral response. Other studies have shown the risk ADE poses, as well as potential strategies to mitigate it. However, as the interaction between DENV and the adaptive immune system is continued to be mapped, the engineering gap to actually create an effective and safe DENV vaccine is not necessarily clear. Here, we have taken an important design leap towards establishing a pan-DENV vaccine by synthesizing the first mRNA-LNP vaccines that drives potently neutralizing antibody responses against all DENV serotypes. We have successfully generated monovalent mRNA vaccines that induce protective immunity against all four DENV serotypes in monovalent and tetravalent formulations. Further we have demonstrated that deletion of the fusion loop epitope can dampen antibody dependent enhancement, thereby increasing vaccine safety. These preliminary data establish the feasibility of the mRNA vaccine encoding for prM/ENV as a viable vaccination strategy that supports the further development and optimization of this platform.

## MATERIALS AND METHODS

### Viruses and Cells

DENV serotype 1 16007 and DENV 2 serotype New Guinea C (NGC) were provided by Michael S. Diamond at Washington University in St. Louis, MO. DENV 3 serotype H87 and DENV 4 serotype UIS 497 were obtained through BEI resources (VR-80 and NR-49724 respectively) NIAID, NIH, as part of the WRCEVA program. All DENV serotypes were cultured in C6/36 cells at 28C and titer determined by focus forming assay (FFA). Several detection monoclonal antibodies used in FFA’s, including 9.F.10 (Santa Cruz Biotech, cat: SC-70959) and 4G2 (BEI resources, NR-50327). All DENV experiments were conducted under BSL-2 (BSL-2) conditions at either the University of Illinois Chicago (UIC) College of Medicine, St. Louis University, or the University of Kentucky. Vero E6 (CRL 1586) and K562 (CCL-243) cells were obtained from American Type Culture Collection (ATCC) and maintained per provided guidelines. HEK 293T cells were provided by Donna MacDuff at UIC.

### mRNA synthesis and LNP encapsulation

Constructs were generated encoding for the tPA signal sequence upstream of the native DENV2, DENV3, or DENV4 prM/ENV coding regions. Strains selected were: DENV2 (strain Thailand 16684/84), DENV 3 (strain Sri Lanka D3/H/IMISSA/SRI/2000/1269), and DENV 4 (strain Dominica/ 814669/1981). The stem/transmembrane domains of envelope proteins were replaced with the JEV stem/transmembrane domain, and the fusion loop epitope was mutated with the three amino acid mutations (G106R, L107D, and F108A). The sequence of constructs is provided in Supplemental Material. Source DNA plasmids encoding the vaccine mRNA were linearized, and the T7 promoter 5’ of the vaccine sequence was used with T7 *in vitro* transcription (IVT) kits from ThermoFisher/Invitrogen (AM1334 or A57622). All vaccine RNA’s were synthesized with N1 Methylpseudouridine (TriLink Biotech, cat: N-1081). The 5’ cap-1 structures (CellSCript, cat: SCCS2250) and 3’ polyA tails (CellScript, cat: C-PAP5104H) were enzymatically added to all vaccine RNA’s. DENV 2 mRNA’s used only in the immunocompetent monovalent humoral analyses were diluted in Formulation Buffer (Precision Nanosystems, cat: 1002775) were encapsulated into LNP’s using PNI Nanosystems NanoAssemblr using the Genvoy-ILM LNP (Precision Nanosystems, cat: 1002763). All other vaccines were encapsulated into LNP’s using the PNI Ignite with SM102 ionizable lipids (Cayman chemical cat: 35425). All mRNA’s were encapsulated with a 3:1 aqueous:ethanolic ratio at a 12mL/min flow rate. mRNA-LNP’s were concentrated using Amicon Ultra Centrifugal Filter 10kDa MWCO (Milipore Sigma, cat: UFC9019). Encapsulation efficiency of these mRNA-LNP’s were quantified by RiboGreen assay (Invitrogen cat: R11490).

### Mouse experiments

mRNA-LNP vaccines were diluted with PBS to contain their given mRNA dosage (10µg, 20µg, etc) per 50µL and administered intramuscularly in all *in vivo* experiments. 12 week old C57BL/6 mice were purchased from Jackson Laboratory and housed in the pathogen-free Biomedical Resources Laboratory (BRL) at UIC College of Medicine. 8-16-week old C57BL/6 or IFNAR^-/-^ mice were vaccinated intramuscularly in a four-week prime boost manner.

### *In vitro* transfection and electron microscopy

HEK 293T cells were transfected at ∼70% confluence with vaccine mRNA using the Mirus TransIT RNA transfection kit (cat: 2225). Supernatant was collected 24 hours post-transfection and dialyzed overnight at 4C in circulating PBS with a 20,000 molecular-weight-cutoff dialysis cartridge (ThermoFisher cat: 6603). Dialyzed sample was provided to UIC electron microscopy core for imaging using the following parameters. 10-15μl of sample was loaded drop-wise onto a 300-mesh, Formvar/Carbon-coated copper EM grid with excess removed by filter paper via capillary action. One drop 2% Uranyl acetate solution was deposited onto EM grid with excess removed by filter paper via capillary action. Once grid was allowed to dry further, sample was examined via transmission electron microscopy using JEOL JEM-1400F transmission electron microscope, operating at 80 kV. Digital micrographs were acquired using an AMT BioSprint 12M-B CCD Camera and AMT software (Version 701).

### Endpoint Dilution ELISA

Viral stocks of all DENV serotypes were grown at 28C in C6/36 cells for 5-7 days. Supernatant from infected cells was centrifuged at 3,200XG for 10 minutes at 4C to remove cellular debris, then further centrifuged at 141,000XG for 2 hours at 4C with a 20% sucrose cushion. Remaining viral pellets were resuspended in PBS then used to coat ELISA plates. Plates were coated with ∼1E3 FFU’s of purified virus diluted in coating buffer (0.1 M sodium carbonate, 0.1 sodium bicarbonate, 0.02% sodium azide, at pH 9.6) overnight at 4C. Plates were blocked with 2% BSA, .025% Sodium Azide in PBS/T for 1 hour at 37C. All mouse sera was heat inactivated at 56C for 30 minutes before any analysis. Serial dilutions of vaccinated or control mouse sera were added to coated plates and incubated overnight at 4C. Plates were stained with goat anti-mouse IgG-HRP (Invitrogen cat: A16072) for 1 hour at room temperature. ELISA plates were developed briefly using 3,3′,5,5′-tetramethylbenzidine (TMB) (ThermoFisher cat: 34029) and the reaction was stopped using 2N H2SO4. Absorbance was measured using a BioTek ELISA microplate reader at 450nm. Endpoint dilution was calculated by the concentration of sera with signal 2X background level.

### Serum neutralization assay

Focus Reduction Neutralization Tests (FRNT) were performed as described previously.^28,43^ Serial dilutions of vaccinated mouse sera was opsonized with ∼70 FFU’s of a single DENV serotype for 1 hour at 37C. Opsonized virus was added to ∼70% confluent monolayer of VeroE6 cells in a 96 well plate in 2% FBS DMEM. After 1 hour of infection, plates were overlaid with 1% w/v methylcellulose in 2%FBS MEM. Plates were incubated at 37C for 48 (DENV 1, 2, and 3) or 65 hours (DENV 4), then fixed with 4% paraformaldehyde (PFA). Plates were stained with a combination of 9.F.10 (500ng/mL), 4G2, and 1-A1D2 in PermWash Buffer (.1% saponin and BSA in PBS). Plates were washed and stained with goat anti-mouse IgG-HRP secondary antibody in PermWash buffer, then resolved with TrueBlue peroxidase (KPL). Resolved plates were imaged on an ImmunoSpot ELISpot plate scanner (Cellular Technology Limited) and foci were counted with Viridot running on RStudio 3.4.1.^44^

### ADE Assay

Serial dilutions of vaccinated or control mouse sera were opsonized with 1 MOI of a single heterotypic DENV serotype (DENV 1 16007, DENV 2 New Guinea C, or DENV 3 H87) for 1 hour at 37C. Opsonized virus was mixed with FcγR+ K562 cells in a 96 well V-bottomed plate. 16 hours post infection, cells were fixed with 4% PFA and stained with a 4G2 chimeric rabbit mAb (Invitrogen cat: MA5-47848) overnight at 4C. Plates were washed and stained with an anti-rabbit fluorescent secondary antibody (Invitrogen cat: A-11008) and analyzed via flow-cytometry.

## Data analysis

Figures and statistical analysis were predominately conducted on GraphPad Prism. Statistical significance was was determined by unpaired one-way ANOVA’s for multiple comparisons in sera analysis. Students T-Test’s were used in ADE assays, and log-rank tests were used for survival curve comparisons. DENV foci were counted with Viridot: An automated virus plaque (immunofocus) counter for the measurement of serological neutralizing responses with application to dengue virus running on RStudio 3.4.1. Flow data was analyzed with BD Biosciences FlowJo. Infographic’s in figures were designed using BioRender.

## ACKNOWLEDGEMENTS

The following reagent was obtained through BEI Resources, NIAID, NIH: Monoclonal Anti-Flavivirus Group Antigen, Clone D1-4G2-4-15 (produced in vitro), NR-50327. This work was funded by NIH R01 AI150672 to Justin Richner, 5P20GM148326-02 to Amelia Pinto, and 3U01CA260541-02S2 to James Brien.

